# Revealing Pentose Catabolism in *Pseudomonas putida*

**DOI:** 10.1101/2022.10.05.510959

**Authors:** Mee-Rye Park, Rahul Gauttam, Bonnie Fong, Yan Chen, Hyun Gyu Lim, Adam M. Feist, Aindrila Mukhopadhay, Christopher J. Petzold, Blake A. Simmons, Steven W. Singer

**Affiliations:** Joint BioEnergy Institute, Emeryville, CA, 94608; Biological Systems and Engineering Division, Lawrence Berkeley National Laboratory, Berkeley, CA, 94720; Department of Bioengineering, University of California San Diego, 9500 Gilman Dr., La Jolla, CA 92093, USA

**Author notes:** **Corresponding author:** Steven Singer, 5885 Hollis Street, Emeryville, CA 94608, http://jbei.org. These authors contributed equally to this work (Mee-Rye Park, Rahul Gauttam). **Data availability statement** The whole genome sequences for five *Pseudomonas* species have been deposited in GenBank under these accession numbers: *Pseudomonas* sp. BP6 (<u>JAGINI000000000</u>), *Pseudomonas* sp. BP7 (<u>JAGINJ000000000</u>), *Pseudomonas* sp. BP8 (<u>JAGINK00000000</u>), *Pseudomonas* sp. M2 (<u>JADOUD010000001</u>), *Pseudomonas* sp. M5 (<u>JAFBBH000000000</u>). The mass spectrometry data have been deposited to the ProteomeXchange Consortium via the PRIDE partner repository with the dataset identifier PXD031549 and 10.6019/PXD031549. **CRediT (Contribution Roles Taxonomy)** M.-R.P. and S.W.S designed project; M.-R. P., R. G., B.F., Y.C. performed experiments; H.G.L. and A.M.F. obtained sequencing data; A.M., C.J., B.A.S. and S.W.S acquired funding and supervised research; M.-R.P., R.G. and S.W.S wrote the manuscript and all authors approved.

## Abstract

The *Pseudomonas putida* group in the Gammaproteobacteria has been intensively studied for bioremediation and plant growth promotion. Members of this group have recently emerged as promising hosts to convert intermediates derived from plant biomass to biofuels and biochemicals. However, most strains of *P. putida* cannot metabolize pentose sugars derived from hemicellulose. Here we describe three isolates that provide a broader view of the pentose sugar catabolism in the *P. putida* group. One of these isolates clusters with the well-characterized *P. alloputida* KT2440 (strain BP6); the second isolate clustered with plant growth-promoting strain *P. putida* W619 (strain M2), while the third isolate represents a new species in the group (strain BP8). Each of these isolates possessed homologous genes for oxidative xylose catabolism (*xylDXA*) and a potential xylonate transporter. Strain M2 grew on arabinose and had genes for oxidative arabinose catabolism (*araDXA*). A CRISPRi system was developed for strain M2 and identified conditionally essential genes for xylose growth. A glucose dehydrogenase was found to be responsible for initial oxidation of xylose and arabinose in strain M2. These isolates have illuminated inherent diversity in pentose catabolism in the *P. putida* group and may provide alternative hosts for biomass conversion.

**Originality-Significance Statement:** Members of the *Pseudomonas putida* group are intensively studied for their role in plant growth promotion and biomass conversion. Despite this interest, the scope of pentose oxidation, key sugars in plant biomass, in this group is not known. Here, we report targeted isolation of members of the *P. putida* group that grow by xylose and arabinose oxidation. Using a combined genomic and proteomic approach, we identify gene products involved in pentose oxidation and identify conditionally essential genes for xylose oxidation using a CRISPRi gene repression approach. This work describes a targeted isolation and analysis strategy that may applied for many microbial groups of industrial and agricultural interest.

## INTRODUCTION

Lignocellulosic biomass is an abundant and sustainable global source as feedstocks for the production of biofuels and bio-based products (Paul and Dutta, 2018; Dahmen et al., 2019). Biofuels and bio-based chemicals have been traditionally produced from lignocellulosic hydrolysates by microorganisms such as *Saccharomyces cerevisiae* and *Escherichia coli* (Liu et al., 2021). However, these microbial hosts are limited in substrate range and are sensitive to toxic inhibitors that are often present in hydrolysates (Piotrowski et al., 2014). Therefore, other potential hosts with broader substrate ranges and higher tolerance to inhibitors have been developed to complement *S. cerevisiae* and *E. coli* (Keasling et al., 2021). Among the most promising is *Pseudomonas putida* KT2440, which has been recently reclassified as *Pseudomonas alloputida* KT2440 (Shi et al., 2017; Shields-Menard et al., 2018; Wang et al., 2018; Dong et al., 2019; Bentley et al., 2020). *P. alloputida* is a representative of the *P. putida* group in the Gammaproteobacteria, members of which have been intensively studied for their role in bioremediation and plant growth promotion. *P. alloputida* KT2440 is of particular interest because of its ability to convert plant-derived aromatics, and has been engineered to produce a variety of biofuels and bio-based chemicals from both glucose and aromatics (Dong et al., 2019; Johnson et al., 2019; Banerjee et al., 2020).

Pentose sugars xylose and arabinose are the predominant constituents of hemicellulose; xylose makes up a substantial amount of the total plant sugars (10-25% of dry biomass) followed by arabinose (usually 2-3%, although some hydrolysates contain up to 20%) (Zhang et al., 2014; Agrawal et al., 2015; Rocha et al., 2015; Kumar et al., 2018; Dehghanzad et al., 2020; Narisetty et al., 2022). However, *P. alloputida* KT2440 is not able to catabolize pentose sugars (Isikgor and Becer, 2015; Lim et al., 2021). Therefore, expanding the substrate range of *P. alloptuida* to include pentose sugars will improve the overall carbon conversion efficiency of lignocellulosic hydrolysates. Several approaches have been used to engineer *P. alloputida* to utilize pentose sugars. The xylose isomerase pathway from *E. coli*, which convers xylose to intermediates in the pentose phosphate pathway, has been expressed in *P. alloputida* and used to convert xylose to *cis-cis*-muconic acid (Ling et al., 2022). Two oxidative pathways for xylose catabolism that proceed through xylonate as an intermediate, the Weimberg pathway, originally characterized in *Caulobacter crescentus*, and the Dahms pathway from *E. coli*, have been expressed in *P. alloputida* and have been used to produce rhamnolipids and indigoidine (Bator et al., 2020; Lim et al., 2021). Heterologous expression of isomerase (*E. coli*) and oxidative (*Burkholderia ambifaria*) pathways for arabinose catabolism have allowed it to grow on this pentose sugar (Elmore et al., 2020). However, problems such as genetic instability, long lag-phases and low cell density have been encountered during these engineering efforts (Jeffries, 2006; Meijnen et al., 2009; Kang et al., 2018; Elmore et al., 2020).

A complementary approach to genetic engineering is to obtain isolates related to *P. alloputida* KT2440 that can natively grow on pentose sugars. These isolates may serve as alternative hosts for the production of biofuels and bio-based chemicals. One member of the *P. putida* group, *Pseudomonas taiwanensis*, has been shown to grow on xylose (Köhler et al., 2015). Analysis of the *P. tawianensis* genome indicated that it possessed a variant of the oxidative xylose pathway found in *C. crescentus*, converting xylose through oxidative intermediates (xylonate and 2-keto-3-deoxyxylonate) to α-ketoglutarate. The *P. taiwanensis* pathway has been expressed in *P. alloputida* KT2440, conferring on it the ability to grow on xylose (Bator et al., 2020). Here we describe a targeted isolation approach to obtaining *Pseudomonas* species related to *P. alloputida* KT2440 that grow on xylose and arabinose. These isolates illuminate the extent of pentose catabolism in the *P. putida* group and provide possible new hosts for metabolic engineering.

## EXPERIMENTAL PROCEDURES

### Soil collection, microbial isolation, and screening

A total of 40 environmental samples were collected from the diverse habitats in Emeryville, CA, USA to isolate microbes. The samples were collected in sterile zip-lock plastic maintaining aseptic conditions and brought to the laboratory, then stored at 4°C. Culturable bacteria were isolated from liquid suspensions prepared at approximately 2.5% (w/v) in autoclaved minimal medium. Serial dilution of the sample suspensions was plated on *Pseudomonas* isolation agar (PIA) plates (Sigma-Aldrich, St. Louis, MO, USA) first and incubated for 1-2 days at 30 °C. Visible bacterial colonies were selected and were sub-cultured by streaking on M9 agar plates containing 0.5% (w/v) xylose and *p*-coumarate, respectively, in a sequence. For purification, a single bacterial colony was re-streaked on the same medium several times. The growth of each colony was subsequently screened in M9 minimal media supplemented with 0.5% (w/v) glucose, xylose and *p*-coumarate as a sole carbon source at 30 °C. Following the screening of isolation to obtain pure cultures, selected strains were routinely cultured in LB at 30 °C overnight. The pure cultures of the isolates were preserved in 20% glycerol stock at -80 °C.

### Cultivation conditions

All the *Pseudomonas* strains were routinely propagated in Luria Bertani (LB) media (tryptone 10 g/L, yeast extract 5 g/L, and NaCl 2.5 g/L) at 30 □C and 200 rpm. For the growth experiments and sample preparation for omics analysis, the cells were grown in minimal medium containing 6 g/L (Na_2_HPO_4_), 3 g/L KH_2_PO_4_, 1.4 g/L (NH_4_)_2_SO_4_, 0.5 g/L NaCl, 0.2 g/L MgSO_4_.7H_2_O, 0.015 g/L CaCl_2_.H_2_O, and 1 mL/L trace element solution purchased from Teknova (Hollister, CA, USA). Depending on the substrate of interest (glucose, xylose, arabinose, and *p*-coumarate), the minimal medium was supplemented with a carbon source (5 g/L). The seed cultures were prepared by inoculating a single colony from a freshly prepared LB agar plate into the minimal medium supplemented with a carbon source (5 g/L). The next day, seed cultures were used as an inoculum to prepare the first pre-culture in minimal medium with a corresponding carbon source. Following day, the strains were inoculated from pre-culture to the main culture (minimal medium with the corresponding carbon source) to start the growth kinetics experiment in 48 well plates with 250 µL of cell culture in each well and for the sample preparation of multi-omics analysis in 50 mL glass tubes. All cultivations were performed in triplicate. The growth was monitored by measuring optical density at 600 nm (OD_600nm_) using a Synergy plate reader (Biotek Instruments, Inc, Winooski, VT, USA). A maximum specific growth rate (h^-1^) was estimated by calculating the slope of the semi-log plot of OD_600nm_ versus time in the exponential growth phase.

### PacBio Sequencing and data analysis

The soil isolates were grown in 5 mL LB broth overnight at 30 °C for high molecular weight genomic DNA sequencing as previously described (Park et al., 2022). The genome sequences for the soil isolates have been deposited in GenBank (Park et al., 2022).

### Phylogenetic analysis

Phylogenetic analysis and tree reconstruction were performed for each genome of isolates obtained in this study, including representative type strains in *Pseudomonas* strains (Keshavarz-Tohid et al., 2017; Keshavarz-Tohid et al., 2019) with complete or draft genome sequences retrieved from GenBank. *Cellvibrio japonicus* Ueda107T was included as outgroup. Nucleotide sequences were aligned using MUSCLE v3.8.425 (Edgar, 2004). Alignments were used to compute Maximum Likelihood analysis using KBase FastTree2 (Price et al., 2010) with built-in branch support values. The resulting tree with Maximum Likelihood analysis was visualized with iTOL software (Heidelberg, Germany) (Letunic and Bork, 2021). Taxonomic species assignment to all newly assembled and downloaded isolates was performed with tool FastANI (Jain et al., 2018) to compute whole-genome similarity metrics such as Average Nucleotide Identity (ANI) values. Each assembly was mapped to each reference strain genome to find orthologous regions using the Mashmap method (reference), for which the average nucleotide index was then calculated and used for comparison. A 95% ANI cutoff is the most frequently used standard for species demarcation (Jain et al., 2018).

### Proteomics and data analysis

Peptide samples were subjected to standard shotgun proteomic analysis protocol (dx.doi.org/10.17504/protocols.io.buthnwj6). Briefly, twenty (20) μg of peptides were separated on a Sigma–Aldrich Ascentis Peptides 588 ES-C18 column (2.1 mm × 100 mm, 2.7 μm particle size, operated at 60 °C) at a 0.400 mL/min flow rate and eluted with the following gradient: initial condition was 98% solvent A (0.1% formic acid) and 2% solvent B (99.9% acetonitrile, 0.1% formic acid). Solvent B was increased to 35% over 11.5 min, and then increased to 80% over 0.5 min, and held for 1.5min, followed by a ramp back down to 2% B over 0.5 min where it was held for 1 min to re-equilibrate the column to original conditions. The eluted peptides were ionized via OptaMaxTM NG Electrospray Ion Source operating in positive ion mode with source and acquisition parameters detailed in the protocol. The MS raw data were acquired using Thermo Scientific Xcalibur version 4.3.73, and the acquired raw data were converted to .mgf files using RawConverter tool and searched against the three annotated *P. putida* strains protein databases with Mascot search engine version 2.3.02 (Matrix Science). Mascot search results are refined by using Scaffold 5.0. Identified peptides are filtered by a 1% peptide-level false discovery rate. In addition, the false discovery rate at the protein level is calculated, and only the proteins with false discovery rate ≤1% are reported. The mass spectrometry data have been deposited to the ProteomeXchange Consortium via the PRIDE partner repository (Perez-Riverol et al., 2022) with the dataset identifier PXD031549 and 10.6019/PXD031549.

### Constructs and strains for CRISPR interference-based gene repression in M2

For adaptation of CRISPR interference in M2 for gene repression studies, *Streptococcus pasteurianus* dCas9-based pRGPdCas9bad and *Streptococcus pyogenes* spdCas9-based pRGPspdCas9badwere used, which were previously adapted for gene repression in KT2440 (Gauttam et al., 2021). The sgRNAs and phenotypic growth measurements were designed following a previously described strategy (Gauttam et al., 2021). The cloning strategy for generating gene-specific CRIPSRi vectors, the complete list for primers containing the sequence for 20-bp homology sequence for gene targeting (Table S8), and the target sequence (Table S9) for a corresponding gene can be found in the Supplementary Information. The test strains generated in *Pseudomona*s sp. M2 (Table S10) and deposited in the public instance of the JBEI registry (http://public-registry.jbei.org/). The strain PP2M491 carrying pRGPspdCas9-bad (with no targeting sgRNA sequence) was used as control. The recombinant strains were grown in minimal medium supplemented with an appropriate carbon source (0.5% w/v), namely, glucose, xylose and arabinose. The oligonucleotides used in this study were ordered from Integrated DNA Technologies (IDT, San Diego, CA, USA) (Table S8).

## RESULTS

### Isolation, screening, and growth characteristics

A screen was designed to isolate *Pseudomonas* species that grew on xylose and plant-derived aromatics (e.g., *p*-coumarate) (Park et al., 2022). Sequential plate-based screens yielded 40 colonies, of which 5 colonies grew in liquid culture on xylose and *p*-coumarate. The growth of these isolates was characterized in minimal medium supplemented with 0.5% (w/v) glucose, xylose, and *p*-coumarate respectively as a sole carbon source and compared to *P. alloputida* KT2440 and *P. taiwanensis* VLB120 as controls. Five isolates were able to grow on all these substrates (Fig. 1). *P. alloputida* KT2440 showed slightly better growth compared to five isolates and *P. taiwanensis* VLB120 on glucose (Fig. 1A and Table 1). In the presence of xylose, the five isolates showed higher growth rates when compared to *P. taiwanensis* VLB120 and BP8 showed the highest optical density among our five isolates (Fig. 1B and Table 1); as expected, *P. alloputida* KT2440 was unable to grow on xylose. The growth rates of the isolates on *p*-coumarate were comparable to *P. alloputida* KT2440 and *P. taiwanensis* VLB120 showed no growth (Fig. 1C and Table 1). Two isolates (M2 and M5) grew arabinose, whereas the other isolates (BP6-BP8) were not able to grow on arabinose (Fig. 1D and Table 1).

**Figure 1.**
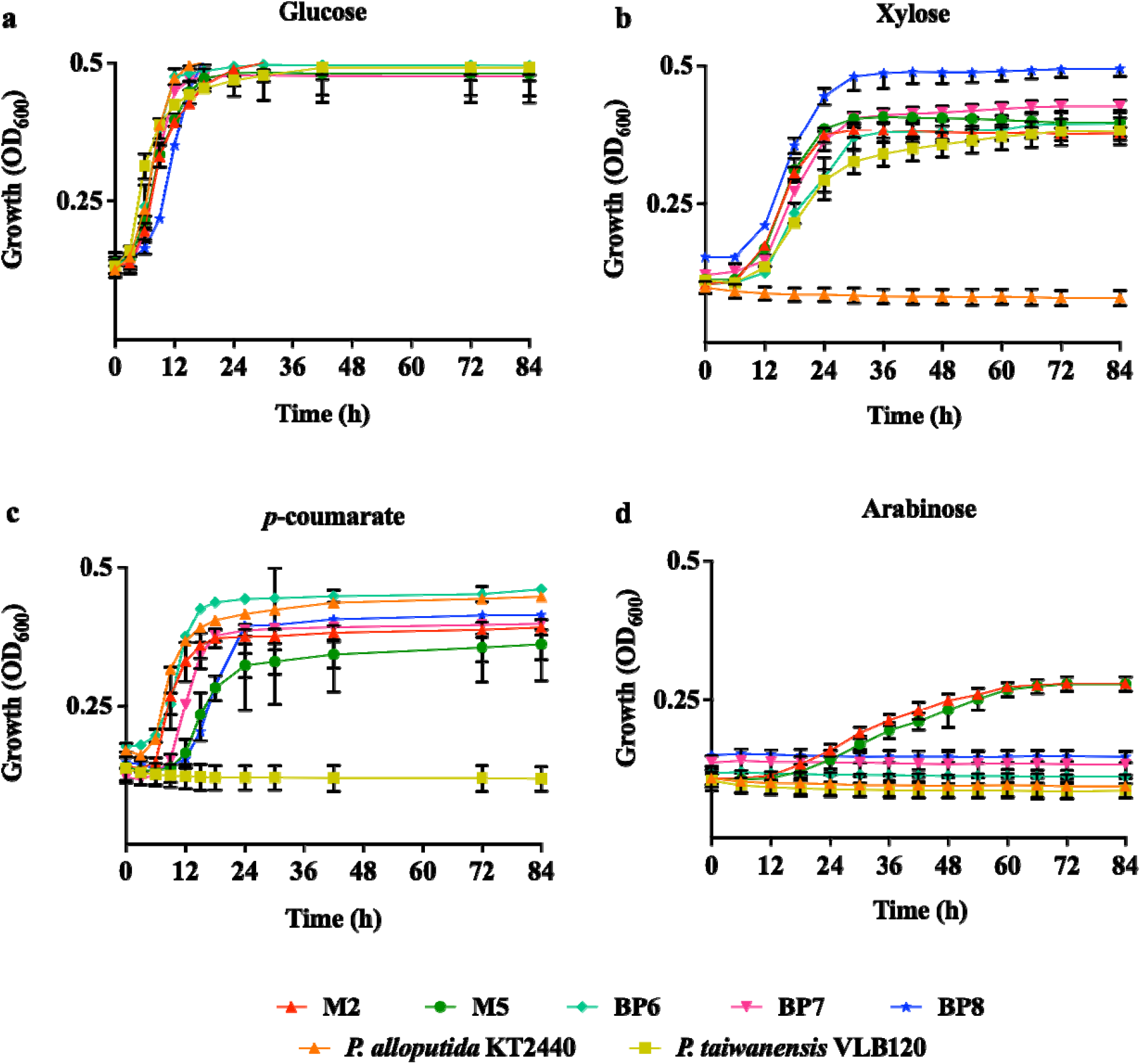
Growth kinetics of five isolates on minimal medium supplemented with 0.5% (w/v) (**a)** glucose, (**b)** xylose, (**c)** *p*-coumarate, and (**d)** arabinose as a sole carbon source. Cell cultures were conducted in biological triplicates and error bars indicate the minimum and maximum values.

**Table 1.** Maximum specific growth rates of five isolates, *P. alloputida* KT2440 and *P. taiwanensis* VLB120 on different carbon sources.

| | Growth rate ($\mu_{\max}$, h<sup>-1</sup>)<sup>a, b</sup> | | | |
| --- | --- | --- | --- | --- |
|  | Glucose | Xylose | Arabinose | *p*-coumarate |
| *P. alloputida* KT2440 | 0.90 ± 0.01 | ND<sup>c</sup> | ND | 0.63 ± 0.10 |
| *P. taiwanensis* VLB120 | 0.77 ± 0.11 | 0.28 ± 0.02 | ND | ND |
| *P. putida* M2 | 0.84 ± 0.03 | 0.46 ± 0.02 | 0.14 ± 0.01 | 0.56 ± 0.01 |
| *P. putida* M5 | 0.70 ± 0.04 | 0.49 ± 0.04 | 0.14 ± 0.02 | 0.35 ± 0.02 |
| *P. alloputida* BP6 | 0.77 ± 0.03 | 0.36 ± 0.04 | ND | 0.56 ± 0.04 |
| *P. alloputida* BP7 | 0.70 ± 0.04 | 0.42 ± 0.04 | ND | 0.49 ± 0.04 |
| *Pseudomonas* sp. BP8 | 0.70 ± 0.02 | 0.49 ± 0.03 | ND | 0.42 ± 0.01 |

<sup>a</sup> Mean value and standard deviation calculated based on three biological replicates.
<sup>b</sup> Growth determined in microtiter plates at 30°C and pH 7. A conversion factor of 7 between a conventional bench-top and a microtiter plate was applied to estimate the growth rate.
<sup>c</sup> ND, not determined due to no growth

### Whole genome-derived phylogenetic classification of the isolates

The phylogenetic affiliation of the five pentose-utilizing isolates was determined by whole-genome comparisons to a representative set of *Pseudomona*s isolates. The five isolates were affiliated with the *P. putida* group and are clearly distinguished from other *Pseudomonas* groups (Fig. 2). Within the *P. putida* group, M2 and M5 were found to have the closest matches to the *P. putida* W619 (96.5% ANI). BP6 and BP7 were found to be closely matched to the species *P. alloputida* LF54 (96.5% ANI). The genome sequences of M2/M5 (99.1% ANI) and BP6/BP7 (99.9% ANI) were nearly identical. BP6/BP7 are the same species as *P. alloputida* KT2440 (96.2% ANI), while M2/M5 are a separate species (86.2% ANI). In contrast, BP8 showed 84.6% ANI to *P. alloputida* KT2440 and does not belong to any neighboring type strains of *Pseudomonas* (Fig. 2). Further, a FastANI analysis with an estimation of ANI represented that BP8 could be considered a new species since the ANI values between BP8 and the closest taxa were much lower than 95% (Table S1), which is the cut-off value for recognizing a new species (Jain et al., 2018).

**Figure 2.**
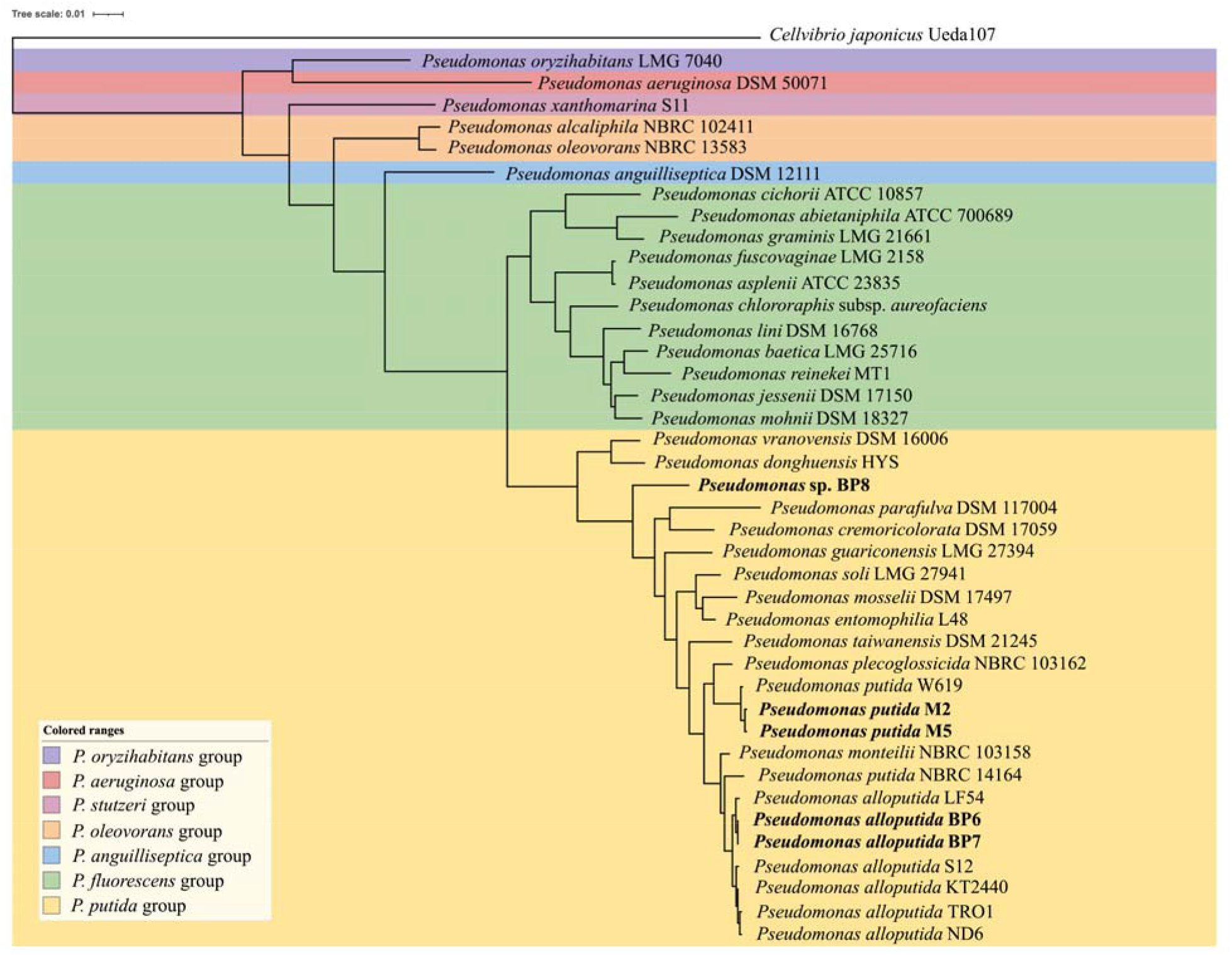
Phylogenetic tree based on whole-genome sequencing of five isolates and *Pseudomonas* type strains. The *Cellvibrio japonicus* Ueda107 sequence was used as the outgroup. The bars indicate sequence divergence. Five isolates in this study are in bold.

As these isolates grew on glucose and *p*-coumarate, the metabolic pathways for these substrates were interrogated in each genome. Strains M2, BP6, and BP8 were chosen as representatives for each phylogenetically distinct pentose-utilizing clade. Each of the isolates contained genes for a complete periplasmic oxidation pathway (*gcd, gnuD* (PP3382-3384), *gnuK, kguK*, and *kguD*), the Entner-Doudoroff pathway (ED; *edd* and *eda*), a phosphorylation protein (*glk*), and the pentose phosphate pathway (PP; *zwfA* and *pgl*); all these enzymes are involved in glucose catabolism (Fig. S1A and S1B). The isolates possess complete pathways for *p*-coumarate catabolism including side-chain oxidation:(*fcs, ech, vdh*, and *pobA*), ring cleavage (*pcaGH, pcaB, pcaC*, and *pcaD*) and the β-ketoadipate conversion (*pcaIJ* and *pcaF*) (Fig. S1A and S1C). The predicted proteins in the glucose catabolism and aromatic degradation pathways are closely related to genes present in *P. alloputida* KT2440 (Nikel et al., 2015; Park et al., 2020).

### Genes for pentose catabolism in *P. putida* isolates

The putative genes for xylose catabolism in the *P. putida* isolates were identified by comparison to the characterized pathway for xylose oxidation in *P. taiwanensis* VLB120. Homologs of xylonate dehydratase (*xylD*), 2-keto-3-deoxyxylonate dehydratase (*xylX*) and α-ketoglutarate dehydrogenase (*xylA*) were present in the M2, BP6 and BP8 genomes, suggesting that the isolates utilize xylose via Weimberg pathway (Fig. 3A-D). Comparing their sequences with the corresponding genes in *P. taiwanensis* VLB120, XylD, XlyX and XylA of three isolates showed 93.8-97.0%, 82.2-90.3%, and 80.3-88.2% identity, respectively (Table S2). In addition, the three genomes and *P. taiwianensis* shared homologs for a LysR-type transcriptional regulator and a permease with three genomes. All the genomes had a second annotated transporter gene (annotated as MHS family metabolite:H+ symporter) between *xylX* and *xylD* and a dehydrogenase related to hydroxypyruvate reductase that were absent in the *P. taiwanenesis* genome.

**Figure 3.**
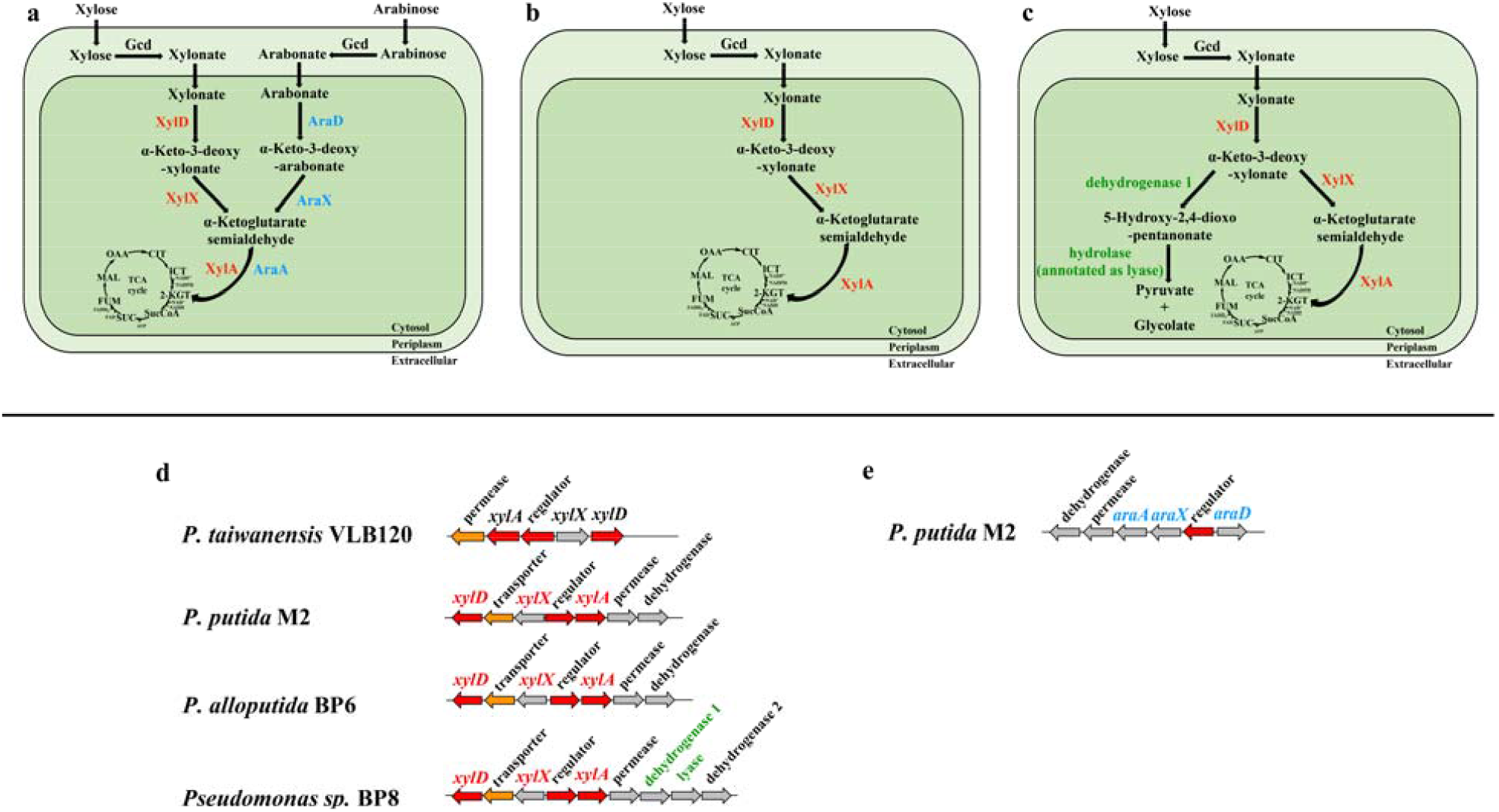
Xylose and arabinose catabolism in three isolates and *P. taiwanensis* VLB120. Schematic overview of xylose and/or arabinose catabolic pathways of **(a)** M2, **(b)** BP6, and **(c)** BP8. Homologous gene clusters are involved in (**d**) xylose and (**e)** arabinose catabolic pathways. Red, orange, and gray arrows indicate cytoplasmic, cytoplasmic membrane, and unknown genes, respectively. Arrow sizes do not represent gene lengths. Abbreviations: OAA, oxaloacetate; CIT, citrate; ICT, isocitrate, 2-KGT, 2-ketoglutarate; SucCoA, succinyl-CoA; SUC, succinate; FUM, fumarate; MAL, malate.

The BP8 genome had an additional annotated dehydrogenase (renamed as dehydrogenase 1 in this study) and lyase adjacent to the putative hydropyruvate reductase (renamed as dehydrogenase 2 in this study), as shown in Fig. 3D. Recent studies reported a non-phosphorylative pathway for xylose catabolism in *Herbaspirillum seropedicae* Z69 (Malán et al., 2021) (identical to *H. seropedicae* SmR1) and *Herbaspirillum huttiense* (Watanabe et al., 2019), respectively, which transform the intermediate of 2-keto-3-deoxy-pentonate (KDP) to 5-hydroxy-2,4-dioxo-pentanonate (HDOP) by a dehydrogenase, and then a HDOP hydrolase is involved in the synthesis of pyruvate and glycolate. In this study, the dehydrogenase 1 and lyase of BP8 are closely related to dehydrogenases (62% identity from both strains) and hydrolases (72% identity from both strains) from Z69 (Malán et al., 2021) and *H. huttiense* (Watanabe et al., 2019). The genomic analysis suggested that BP8 metabolizes xylose in not only the Weimberg pathway but also the non-phosphorylative pathway via the KDP intermediate (Fig. 3C).

Regarding the arabinose catabolism in the M2 genome, genes for an arabonate dehydratase (*araD*) with 43% identity to M2 *xylD*, an α-keto-3-deoxy arabonate dehydratase (*araX*), and a ketoglutarate semialdehyde dehydrogenase (*araA*) were identified that were contained in a cluster (*araD*-*araX*-*araA*) as shown in Fig. 3A and 3E. This gene cluster had an additional gene related to 2-phosphogluconate dehydrogenase.

The sequences of the *xylD* and *araD* genes were used to determine the prevalence of xylose and arabinose oxidation in the *P. putida* group (Fig. 4 and Table S3-4). Interestingly, only a few strains affiliated with the *P. putida* group include the homologs of *xylD* in M2/M5 (Fig. 4). For example, the *xylD* sequence of M2/M5 were >95% similar to the genes of BP6/BP7, *P. putida* W619, *P. alloputida* LF54 and *P. taiwanensis* VLB120, and showed 93% identity with that of BP8 (Table S3). Likewise, when *araD* of M2/M5 was compared against the *Pseudomonas* database, the predicted protein homologs were found in the following 3 species: *P. putida* W619 (100.0% identity), *P. monteilii* NBRC 103158 (98.8% identity) and *P. plecoglossicida* NBRC 103162 (96.2% identity) (Table S4).

**Figure 4.**
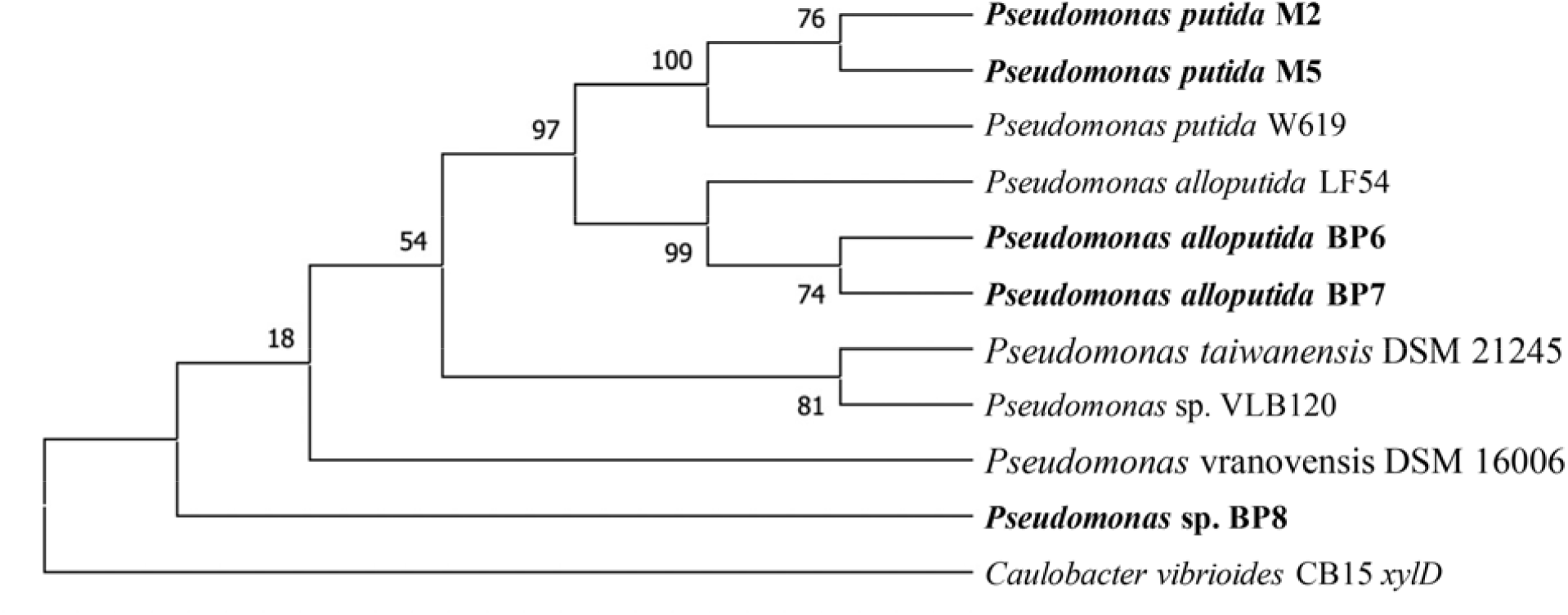
Maximum likelihood trees of xylonate dehydratase gene (*xylD*) from the isolates and closely related proteins (>95% sequence identity threshold) in the *P. putida* group using JTT matrix-based model in MEGA X(MEGA, 2018). The *xylD* of *Caulobacter vibrioides* CB15 was used as the outgroup. Numbers at each node are bootstrap probabilities by 1000 replications. The isolates in this study are in bold.

Comparative proteomics analysis revealed that proteins encoded in the putative operon for xylose oxidation identified by genomic analysis were present at significantly higher abundance during growth on xylose as compared to growth on glucose in the isolates (Table 2). For example, XylD, which converts xylonate to α-keto-3-deoxy-xylonate was present at significantly higher abundance in all three strains (3.7 log_2_ FC in M2; 2.9 log_2_ FC in BP6; 4.6 log_2_ FC in BP8). XylX, which dehydrates α-keto-3-deoxy-xylonate to α-ketoglutarate semialdehyde was also present at significantly higher abundance (5.4 log_2_ FC in M2; 4.5 log_2_ FC in BP6; 5.3 log_2_ FC in BP8), as well as XylA, which reduces α-ketoglutarate semialdehyde dehydrogenase to α-ketoglutarate (4.7 log_2_ FC in M2; 4.3 log_2_ FC in BP6; 5.3 log_2_ FC in BP8). The proteins encoding for the predicted transporter (2.4 log_2_FC in M2; 0.6 log_2_FC in BP6; 2.5 log_2_ FC in BP8) and permease (2.1 log_2_ FC in M2; 1.0 log_2_ FC in BP6; 2.0 log_2_ FC in BP8) in putative xylose oxidation operon were also at higher abundance, suggesting that they also played a role in xylose catabolism. The annotated hydroxypyruvate reductase did not show any evidence of differential abundance. In addition, for strain BP8, the dehydrogenase 1 (4.4 log_2_ FC), the lyase (4.2 log_2_ FC) and the hydroxypyurvate dehydrogenase (dehydrogenase 2) (4.9 log_2_ FC) were present at higher abundance in xylose-grown cells.

**Table 2.**
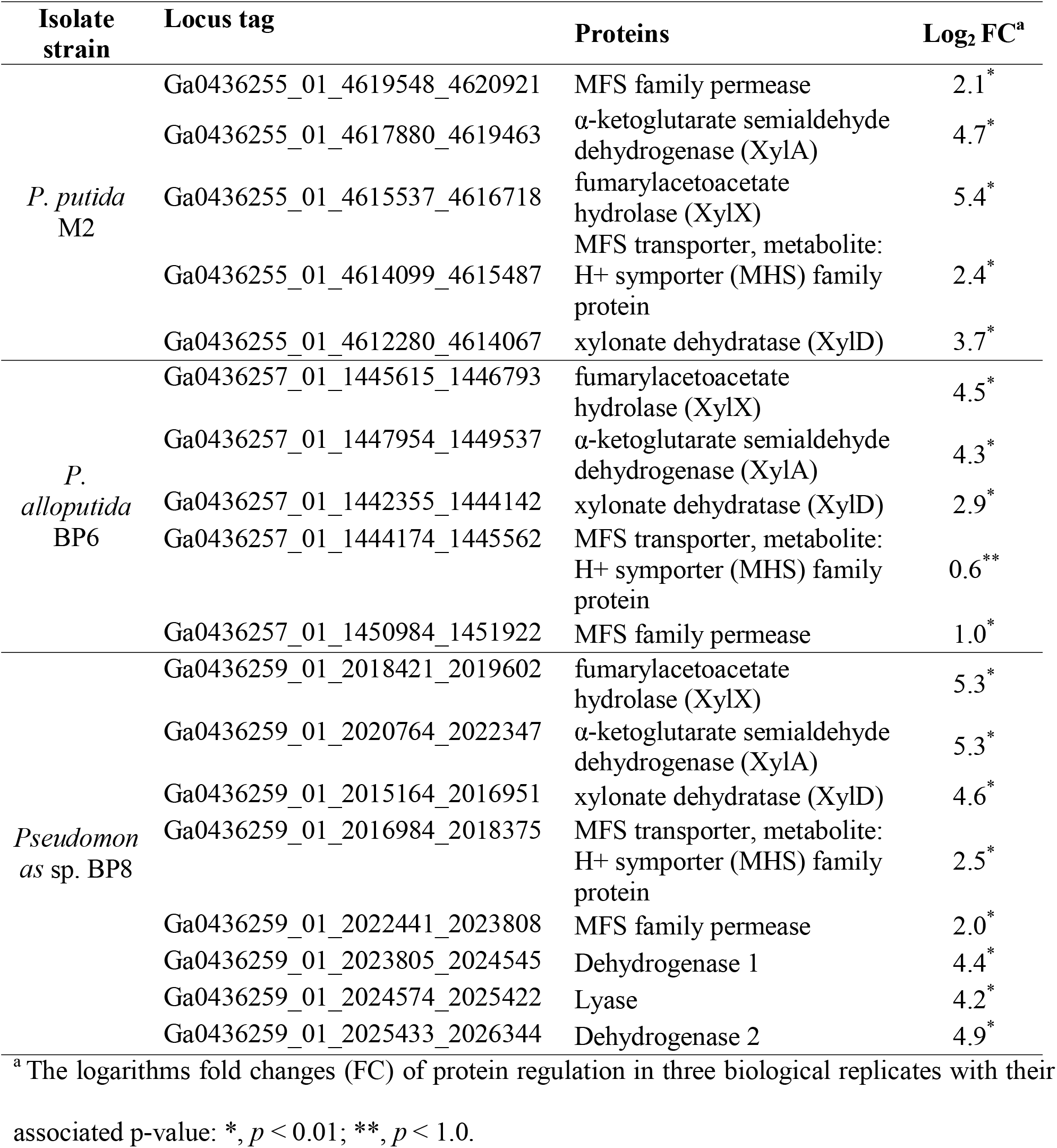
Differential abundance profiles of proteins predicted to be involved in xylose metabolism in M2, BP6 and BP8 grown in 0.5% (w/v) xylose as a sole carbon source compared to glucose in proteomics analysis.

For *P. putida* M2, the arabinose-grown cells had the predicted arabinose oxidation proteins, AraDXA, at higher relative abundance compared to glucose-grown cells (3.4-4.9 log_2_ FC) (Table S5). The permease located between *araX* and *araD*, was also upregulated, suggesting it may be involved in transport of intermediates into the cytosol.

### Identifying conditionally essential genes in strain M2 using CRISPR interference (CRISPRi)

Previously, duet vectors were adapted to express a CRISPRi/dCas9-based gene repression system in *P. alloputida* KT2440 (Gauttam et al., 2021). The study demonstrated the use of two dCas9 homologs from *S. pasteurianus* (dCas9) and *S. pyogenes* (spdCas9) for screening of conditionally essential genes in minimal media. To translate this system to *P. putida*, both dCas9 proteins were used to target six endogenous genes, (acetylglutamate kinase (*argB*), argininosuccinate lyase (*argH*), 2-keto-3-deoxy-6phosphogluconate aldolase (*eda*) phosphogluconate dehydratase (*edd*), chorismate mutase (*pheA*), and orotidine-5’-phosphate decarboxylase (*pyrF*)), which were shown to be essential for strain KT2440 glucose growth in minimal medium. Minimal repression of these conditionally essential genes was observed using the *S. pasteurianus* dCas9-mediated CRISPRi system (Fig. S2). In contrast, growth inhibition was observed for all six genes targeted using the *S. pyogenes* spdCas9-mediated CRISPRi system (Fig. S3).

To demonstrate the utility of the spdCas9-based CRISPRi system in M2, genes shown to be involved in xylose catabolism by proteomics were targeted for gene repression by CRISRPi. (Table S6-9). Repression of these genes did not affect growth on glucose (Fig. S4); however, repression of *xylD* and the predicted transporter located between *xylD* and *xylX* repressed growth on xylose (Fig. 5).

**Figure 5.**
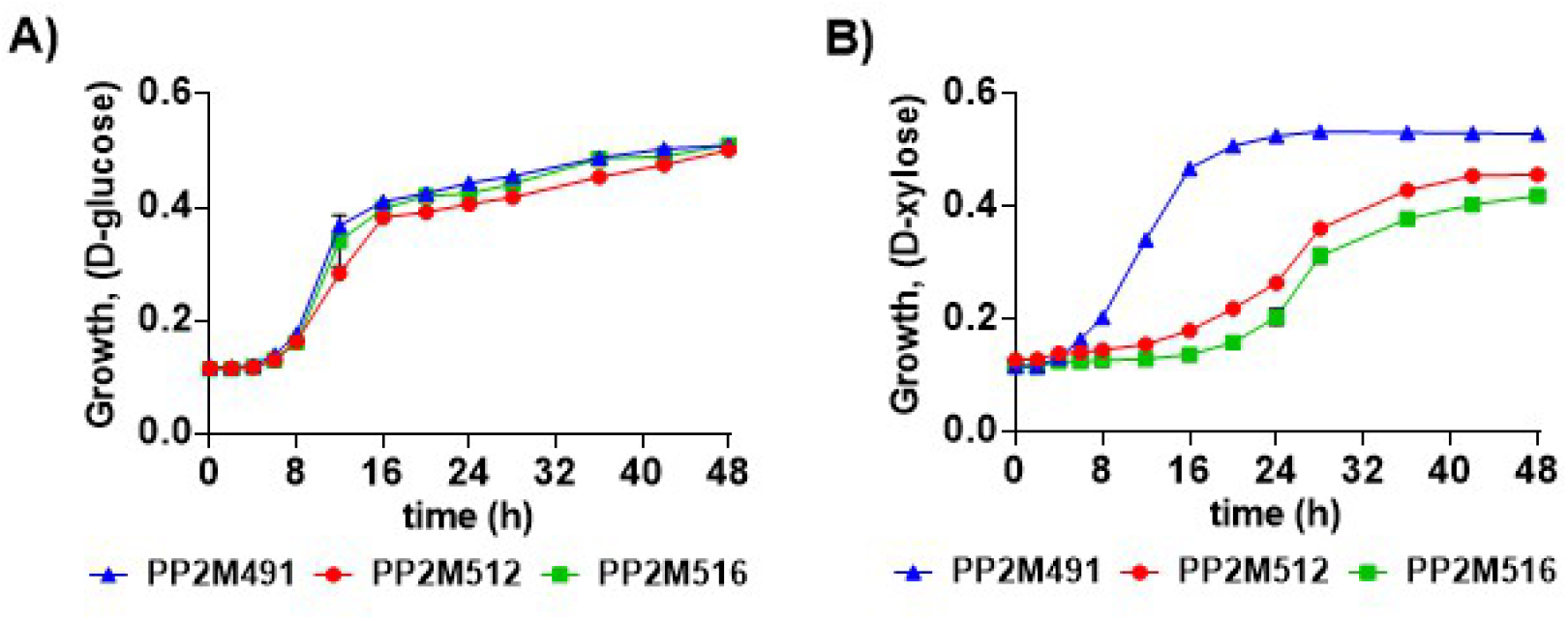
Characterization of putative conditionally essential genes for xylose catabolism in strain M2 using *S. pyogenes* spdCas9-based CRISPRi. The sgRNAs were designed to downregulate the expression of target genes (Gauttam et al., 2021) and sequences are listed in **Table S9**. For strain descriptions, refer to **Table S6**. The growth phenotype was assessed in minimal medium with different C-sources namely, glucose **(A)** and xylose **(B)** for the recombinant strains PP2M512 (pRGPspdCas9bad-*xyl transporter*) and PP2M516 (pRGPspdCas9bad-*xylD*). The growth was compared to control strain PP2M491 (pRGPspdCas9bad) carrying a vector with no targeting sgRNA sequence. Each graph represents the mean values of biological triplicates from at least three individual cultivations, and error bars represent standard deviations.

### Glucose dehydrogenase (Gcd) is responsible for xylose and arabinose oxidation

In comparison to the well-characterized oxidative pentose pathway in *Caulobacter crescentus*, the oxidative pathway in the *Pseudomonas* isolates lacked an evident xylose or arabinose dehydrogenase. The annotated hydroxypyruvate reductase (xylose) and phosphogluconate-2-dehydrogenase (arabinose) from *P. putida* M2 were expressed in *E. coli* but showed minimal activity on xylose and arabinose, suggesting that they were not involved in C5 catabolism (data not shown). Since these dehydrogenases characterized from M2 were not pentose dehydrogenases, the enzymes responsible for xylose and arabinose oxidation remained unknown. A gene coding for glucose dehydrogenase (*gcd*) was targeted for gene repression, based on previous work in *P. taiwanensis* and *P. alloputida*, which demonstrated that xylose oxidation was dependent on Gcd. The recombinant CRISPRi strains were cultured in minimal media supplemented with glucose, xylose or arabinose (0.5% w/v). Repression of *gcd* inhibited strain M2 growth in minimal media with xylose and arabinose as a sole carbon source (Fig. 5B and 5C). However, the growth was not affected on glucose, suggesting *gcd* is not a conditionally essential gene for M2 growth on glucose (Fig. 6).

**Figure 6.**
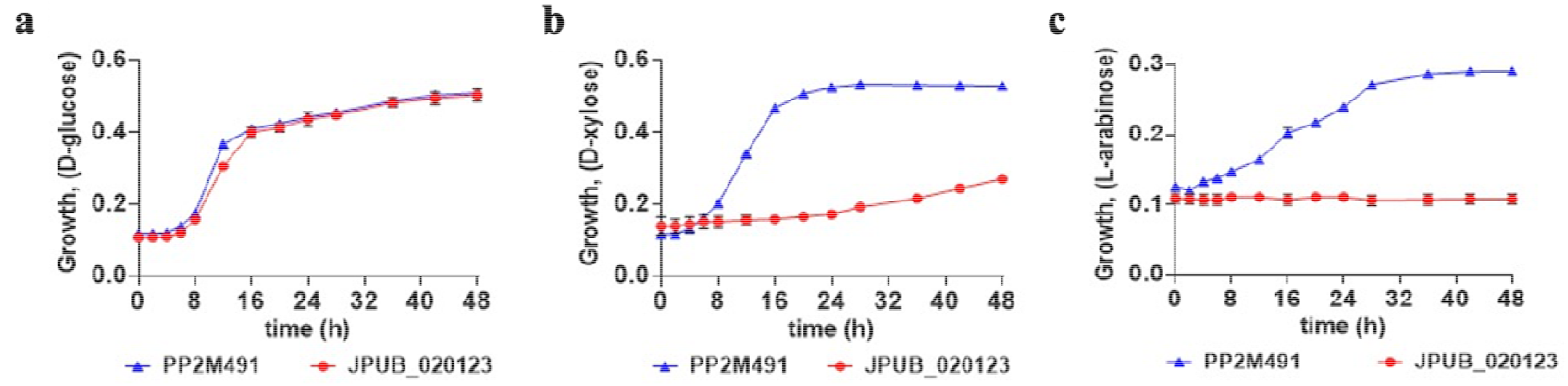
Phenotypic characterization of Gcd using *S. pyogenes* spdCas9-based CRISPR interference in minimal medium using different carbon sources: (**a)** glucose, (**b)** xylose and (**c)** arabinose for recombinant M2 strain JPUB_020123 (pRGPspdCas9bad-*gcd*). The growth was compared to control strain PP2M491 (pRGPspdCas9bad) carrying a vector with no targeting sgRNA sequence. Each graph represents the mean values of biological triplicates from at least three individual cultivations, and error bars represent standard deviations.

## DISCUSSION

This study has provided a broad perspective on the ability of members of the *P. putida* group to metabolize pentose sugars relevant to lignocellulose bioconversion. Targeted isolation studies carried out with xylose and *p*-coumarate as the targeted substrates provided isolates that expressed an oxidative pathway for xylose catabolism similar to the Weimberg pathway in *C. crescentus* (Shen et al., 2020). Comparison of the isolates to the *P. putida* group genomes indicates that xylose oxidation is relatively restricted in this group. Strain BP6 and BP7 cluster with *P. alloputida*, the species which is the most thoroughly characterized of the *P. putida* strains, including strain KT2440 (Keshavarz-Tohid et al., 2019). The BP6 and BP7 strains are most closely related to *P. alloputida* LF54, which is a representative of the most divergent clade of *P. alloputida*, (Passarelli-Araujo et al., 2021). The M2 and M5 strains are affiliated with *P. putida* W619, which has been characterized for its ability to promote the growth of plants (Taghavi et al., 2005). *P putida* W619 has been shown to grow on both xylose and arabinose (Davis et al., 2013). The survey also revealed other members of the *P. putida* group (*Pseudomonas sp*. BP8 and *P. vranovensis*) that grew on xylose along with the previously characterized *P. taiwanensis*. Interestingly, the ability to grow on arabinose is even more constrained, and is only present in *P. monteilii* and *P. plecoglossicida* along with the clade containing *P. putida* M2/M5 and W619. These results are consistent with pentose oxidation being a niche activity in the *P. putida* group. Strain BP8 had genes for additional pathway for xylose oxidation that may divert intermediates in the oxidative pathway to pyruvate and glycolate. Proteins for both pathways were present in the xylose-grown cells and that may have contributing to the higher growth rate of BP8 relative to the strains with one pathway.

Integrated genomic and proteomic analyses demonstrated that homologs of the Weimberg pathway (XylDXA and AraDXA) were responsible for xylose and arabinose oxidation in the *P. putida* group strains. These proteins are likely responsible for converting the oxidized sugar (xylonate and arabinoate) through multiple dehydrations and an oxidation to produce α-ketoglutarate. Repression of the genes encoding these proteins using CRISPRi in *P. putida* M2 demonstrated that only *xylD* interference repressed xylose growth. This result is consistent with studies in *P. alloputida* KT2440 that expression of only *xylD* is required to confer growth on xylose by these isolates, and that the activities of XylX and XylA can be recruited using other proteins (Lim et al., 2021). In the putative operon for xylonate oxidation, there is a transcriptional regulator and permease found in common in all the strains. However, the newly isolated strains have an annotated transporter (MHS family metabolite:H+ symporter) that is not present in *P. tawianensis*. The presence of the transporter, which was identified in the proteomes of strains M2, BP6 and BP8, may improve growth (0.36-0.49 h^-1^) on xylose relative to *P. taiwanensis* VLB120 (0.28 h^-1^). The importance of this transporter was reinforced in the *P. putida* M2 CRISPRi experiments, as repression of this transporter inhibited the xylose growth of strain M2. The possible substrate for the transporter is xylonate, as both CRISPRi-based repression of *gcd* in strain M2 and deletion of *gcd* in strain BP6 affected the ability of the strains to grow on xylose, indicating that Gcd oxidizes xylose to xylonate in the periplasm and then it is transported into the cytosol by the transporter present in the isolates described here. The xylose dehydrogenase activity of Gcd is consistent with previous studies on *P. alloputida* KT2440 (Meijnen et al., 2009), *P. putida* NCTC 10936 (Hardy et al., 1993) and *P. taiwanensis* VLB120 (Köhler et al., 2015), which demonstrated that Gcd was required for xylose oxidation. Interestingly, CRISPRi experiments also demonstrated that Gcd was required for arabinose growth, suggesting that arabinose is also oxidized to arabinoate in the periplasm and then the arabinoate is transported into the cytosol. The requirement of Gcd for pentose-based growth in multiple *P. putida* group strains and for multiple substrates indicates there is a link in the metabolism of all three major lignocellulose-derived sugars in the members of the *P. putida* group. The catalytic promiscuity of Gcd for lignocellulose-derived hexose and pentose sugars has also been demonstrated in sugar catabolism in *Sulfolobus solfataricus* and *Sulfolobus acidocaldarius* (Nunn et al., 2010).

The growth rates of the native xylose-oxidizing strains in the *P. putida* group are comparable or better than strains of *P. alloputida* KT2440 that have been engineered to grow on xylose (Elmore et al., 2022). Further, use of adaptive laboratory evolution with the strains characterized here will likely result in additional diversity and useful strain-specific catabolic properties. Therefore, these isolates may be suitable complements to strain KT2440 for metabolic engineering focused on lignocellulose bioconversion.

## ACKNOWLEDGEMENTS

Dr. Birgitta Ebert (University of Queensland) is acknowledged for providing *Pseudomonas taiwanensis* VLB120.

