## Supplemental Information for "Revealing Pentose Catabolism in *Pseudomonas putida*"

http://jbei.org

**Supplementary Figures**

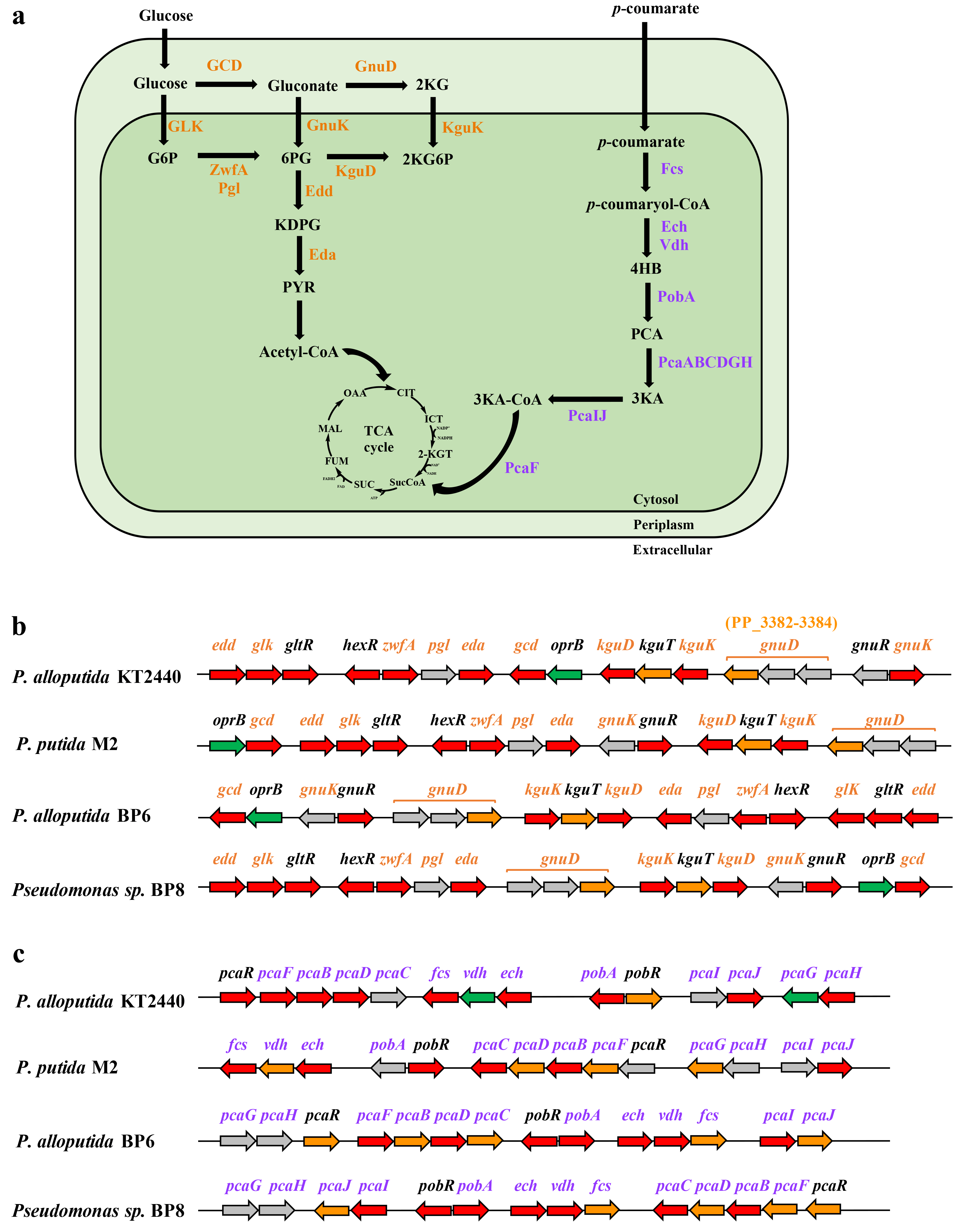

**Figure S1.** (**a)** Schematic overview of glucose catabolism and *p*-coumarate degradation in the isolates and *P. alloputida* KT2440. Gene organization of clusters of (**b)** peripheral pathway, pentose phosphate pathway, Entner-Doudoroff (ED) pathway and Embden-Meyerhof-Parnas (EMP) pathway involved in glucose catabolism; (**c)** *p*-coumarate degradation and *β*-ketoadipate pathways. Red, orange, green and gray arrows indicate cytoplasmic, cytoplasmic membrane, outer membrane, unknown genes, respectively. Arrow sizes do not represent gene lengths. The abbreviations used are: 2-KG, 2-ketogluconate; G6P, glucose-6-phosphate (P); 6PG, 6-phosphogluconate; 2KG6P, 2-ketogluconate-6-P; KDPG, 2-dehydro-3-deoxy-phosphogluconate; 4HB, 4-hydroxybenzoate; PCA, protocatechuate; 3KA, 3-ketoadipate; 3KA-CoA, 3-ketoadipyl-CoA; Pyr, pyruvate; OAA, oxaloacetate; CIT, citrate; ICT, isocitrate, 2-KGT, 2-ketoglutarate; SucCoA, succinyl-CoA; SUC, succinate; FUM, fumarate; MAL, malate.

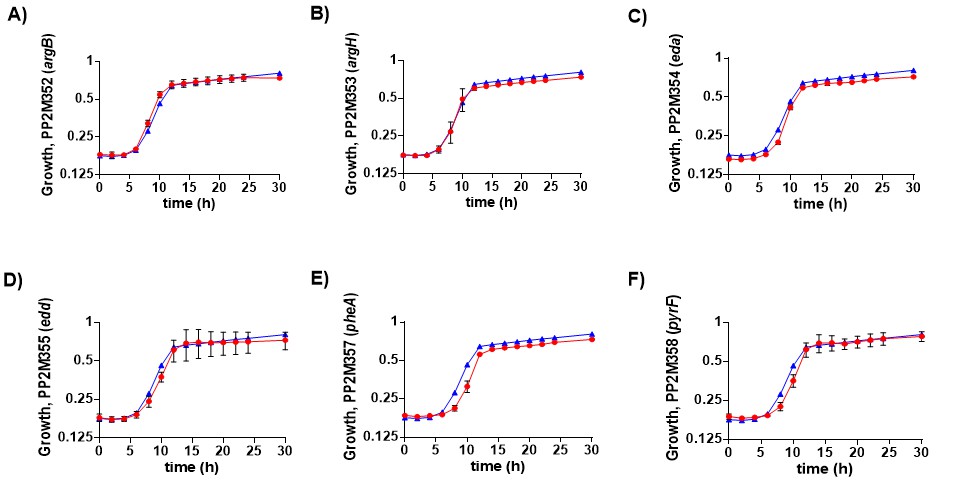

**Figure S2.** Characterization of putative conditionally essential genes in new host M2 using *S. pasteurianus* dCas9 based CRISPRi **(A-E)**. sgRNAs were designed to downregulate the expression of target genes using the details from our previous study (Gauttam et al., 2021) and sequences are listed in **Table S9**. For strains description, refers to **Table S6**. Growth phenotype was assessed in minimal medium with glucose (0.5% w/v) as C-source supplemented with inducers IPTG (1 mM) and arabinose (0.2% w/v) for the recombinant strains PP2M352 (pRGPdCas9bad-*argB*) PP2M353 (pRGPdCas9bad-*argH*), PP2M354 (pRGPdCas9bad-*eda*), PP2M355 (pRGPdCas9bad-*edd*), PP2M357 (pRGPdCas9bad-*pheA*) and PP2M358 (pRGPdCas9bad-*pyrF*). The growth was compared to control strain PP2M350 (pRGPdCas9bad) carrying vector with no targeting sgRNA sequence. In each graph the control strain PP2M350 is represented as a blue triangle. The red circle in each graph represents the test strain with downregulation of respective target gene. Each graph represents the mean values of biological triplicates from at least three individual cultivations, and error bars represent standard deviations.

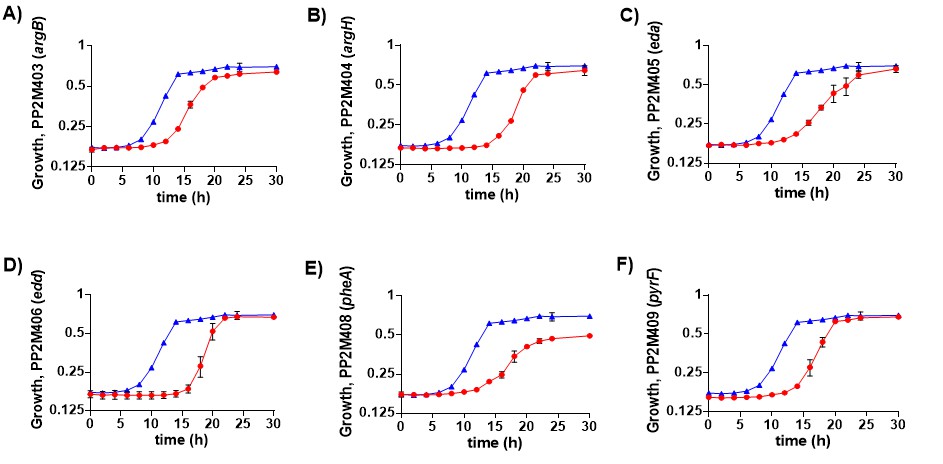
**Figure S3.** Characterization of putative conditionally essential genes in new host M2 using *S. pyogenes* spdCas9 based CRISPRi **(A-F)**. sgRNAs were designed to downregulate the expression of target genes using the details from our previous study (Gauttam et al., 2021) and sequences are listed in **Table S9**. For strain description, refer to **Table S6**. Growth phenotype was assessed in minimal medium with glucose (0.5% w/v) as C-source supplemented with inducers IPTG (1 mM) and arabinose (0.2% w/v) for the recombinant strains PP2M403 (pRGPspdCas9bad-*argB*) PP2M404 (pRGPspdCas9bad-*argH*), PP2M405 (pRGPspdCas9bad-*eda*), PP2M406 (pRGPspdCas9bad-*edd*), PP2M408 (pRGPspdCas9bad-*pheA*) and PP2M409 (pRGPspdCas9bad-*pyrF*). The growth was compared to control strain PP2M491 (pRGPspdCas9bad) carrying a vector with no targeting sgRNA sequence. In each graph the control strain PP2M491 is represented as a blue triangle. The red circle in each graph represents the test strain with downregulation of the target gene. Each graph represents the mean values of biological triplicates from at least three individual cultivations, and error bars represent standard deviations.

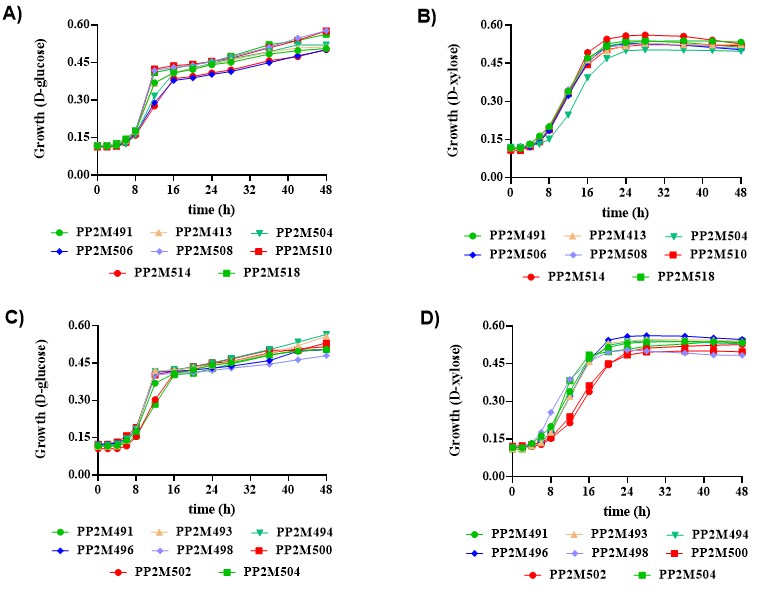

**Figure S4.** Growth phenotype was assessed using *S. pyogenes* spdCas9 for representative genes of pentose sugar (xylose and arabinose) utilization pathway in minimal medium with glucose **(A, C)**, and Xylose **(B, D)**, as C-source. Each graph represents the mean values of biological triplicates from at least three individual cultivations, and error bars represent standard deviations. The description of strains is mentioned in **Table S6**.

**Supplementary Tables**

**Table S1**. Closely related taxon of *Pseudomonas* sp. BP8 with FastANI identity scores in NCBI database

| **Query** | **Closely related taxa** | **ANI %** | **Matches** | **Total** |
| --- | --- | --- | --- | --- |
| *Pseudomonas* sp. BP8 | *Pseudomonas* *entomophila* L48 | 85.5 | 1257 | 2001 |
|  | *Pseudomonas* *plecoglossicida* NBRC 103162 | 85.4 | 1192 | 2001 |
|  | *Pseudomonas* *mosselii* DSM 17497 | 85.4 | 1259 | 2001 |
|  | *Pseudomonas* *soli* LMG 27941 | 85.1 | 1251 | 2001 |
|  | *Pseudomonas* *putida* NBRC 14164 | 85.0 | 1353 | 2001 |
|  | *Pseudomonas* *monteilii* NBRC 103158 | 84.9 | 1305 | 2001 |
|  | *Pseudomonas* *alloputida* ND6 | 84.7 | 1305 | 2001 |
|  | *Pseudomonas* *putida* W619 | 84.6 | 1285 | 2001 |
|  | *Pseudomonas* *alloputida* TRO1 | 84.5 | 1236 | 2001 |
|  | *Pseudomonas* *alloputida* KT2440 | 84.5 | 1285 | 2001 |
|  | *Pseudomonas* *alloputida* S12 | 84.5 | 1295 | 2001 |
|  | *Pseudomonas* *guariconensis* LMG 27394 | 84.4 | 1185 | 2001 |
|  | *Pseudomonas* *alloputida* LF54 | 84.3 | 1240 | 2001 |
|  | *Pseudomonas* *taiwanensis* DSM 21245 | 84.3 | 1232 | 2001 |
|  | *Pseudomonas* *parafulva* DSM 117004 | 84.0 | 1138 | 2001 |
|  | *Pseudomonas* *donghuensis* HYS | 83.7 | 1117 | 2001 |
|  | *Pseudomonas* *cremoricolorata* DSM 17059 | 83.3 | 1062 | 2001 |
|  | *Pseudomonas* *vranovensis* DSM 16006 | 83.0 | 1084 | 2001 |
|  | *Pseudomonas* *rhizosphaerae* DSM 16299 | 81.2 | 794 | 2001 |

**Table S2.** Similarity of y XylDXA proteins of *Pseudomonas taiwanensis* VLB120 with the *Pseudomonas* isolates

|  | **Gene product** | ***Pseudomonas putida* M2** | ***Pseudomonas alloputida* BP6** | ***Pseudomonas* sp. BP8** |
| --- | --- | --- | --- | --- |
| XylD | xylonate dehydratase | 96.6% | 97.0% | 93.8% |
| XylX | fumarylacetoacetate hydrolase | 89.0% | 90.3% | 82.2% |
| XlyA | α-ketoglutarate semialdehyde dehydrogenase | 88.2% | 86.7% | 80.3% |

**Table S3:** Homologous genes of *xylD* of M2 as query sequence in *Pseudomonas* groups.

| **Gene ID** | **Genome** | **Identity** | **Align length** |
| --- | --- | --- | --- |
| Ga0436255_01_4612280_4614067 | *Pseudomonas* *putida* M2 | 100.0% | 595 (100.0%) |
| Ga0436256_01_1387624_1389411 | *Pseudomonas* *putida* M5 | 100.0% | 595 (100.0%) |
| PPUTW619_RS15740 | *Pseudomonas putida* W619 | 99.8% | 595 (100.0%) |
| Ga0436257_01_1442355_1444142 | *Pseudomonas alloputida* BP6 | 98.5% | 595 (100.0%) |
| Ga0436258_01_271208_272995 | *Pseudomonas alloputida* BP7 | 98.5% | 595 (100.0%) |
| O999_RS51000 | *Pseudomonas alloputida* LF54 | 98.3% | 595 (100.0%) |
| H620_RS0111905 | *Pseudomonas taiwanensis* DSM 21245 | 96.8% | 595 (100.0%) |
| PVLB_RS18630 | *Pseudomonas taiwanensis* VLB120 | 96.6% | 595 (100.0%) |
| Ga0436259_01_2015164_2016951 | *Pseudomonas* sp. BP8 | 93.60% | 595 (100.0%) |
| H621_RS0118655 | *Pseudomonas vranovensis* DSM 16006 | 93.6% | 595 (100.0%) |
| BVK86_RS02530 | *Pseudomonas reinekei* MT1 | 92.9% | 595 (100.0%) |
| BLV61_RS03480 | *Pseudomonas mohnii* DSM 18327 | 92.6% | 595 (100.0%) |
| BBI10_RS10270 | *Pseudomonas graminis* LMG 21661 | 89.1% | 595 (100.0%) |
| BLS87_RS25505 | *Pseudomonas abietaniphila* ATCC 700689 | 88.7% | 595 (100.0%) |
| UM91_RS16255 | *Pseudomonas* *oryzihabitans* LMG 7040 | 89.5% | 590 (99.2%) |
| BLW67_RS26765 | *Pseudomonas fuscovaginae* LMG 2158 | 86.7% | 595 (100.0%) |
| BLU37_RS15270 | *Pseudomonas* *asplenii* ATCC 23835 | 86.7% | 595 (100.0%) |

Table S4**:** Homologous genes of *araD* of M2 as a query sequence in *Pseudomonas* groups.

| **Gene ID** | **Genome** | **Ident%** | **Align length** |
| --- | --- | --- | --- |
| PPUTW619_RS10130 | *Pseudomonas putida* W619 | 100.0% | 577 (100.0%) |
| Ga0436255_01_320995_322728 | *Pseudomonas putida* M2 | 100.0% | 577 (100.0%) |
| Ga0436256_01_306509_308242 | *Pseudomonas putida* M5 | 99.8% | 577 (100.0%) |
| Q381_RS0128405 | *Pseudomonas monteilii* NBRC 103158 | 98.8% | 577 (100.0%) |
| PPL01S_RS22130 | *Pseudomonas plecoglossicida* NBRC 103162 | 96.2% | 577 (100.0%) |

**Table S5.** Differential expression profiles of M2 grown in 0.5% (w/v) xylose and arabinose as a sole carbon source in proteomics analysis.

|  | **Locus tag** | **Gene product** | **Log_2_ FC^a^** | |
| --- | --- | --- | --- | --- |
|  |  |  | **Xylose** | **Arabinose** |
| *xyl* operon | Ga0436255_01_4619548_4620921 | MFS family permease | 2.1^*^ | 0.0^*^ |
|  | Ga0436255_01_4617880_4619463 | α-ketoglutarate semialdehyde dehydrogenase (XylA) | 4.7^*^ | -0.5^*^ |
|  | Ga0436255_01_4615537_4616718 | fumarylacetoacetate hydrolase (XylX) | 5.4^*^ | 0.0^*^ |
|  | Ga0436255_01_4614099_4615487 | MHS family metabolite:H+ symporter | 2.4^*^ | 0.0^*^ |
|  | Ga0436255_01_4612280_4614067 | xylonate dehydratase (XylD) | 3.7^*^ | 0.0^*^ |
| *ara* operon | Ga0436255_01_315323_316654 | MFS family permease | 0.0^*^ | 1.1^*^ |
|  | Ga0436255_01_316743_318317 | α-ketoglutarate semialdehyde dehydrogenase (AraA) | 0.2^*^ | 4.9^*^ |
|  | Ga0436255_01_318483_319472 | α-keto-3-deoxy arabonate dehydratase (AraX) | -0.8^*^ | 3.4^*^ |
|  | Ga0436255_01_319501_320808 | MFS family permease | 0.0^*^ | 1.9^*^ |
|  | Ga0436255_01_320995_322728 | arabonate dehydratase (AraD) | 0.3^*^ | 4.7^*^ |

^a^ The logarithms fold changes (FC) of protein regulation in three biological replicates with their associated p-value: *, *p* < 0.01.

**Table S6.** CRISPRi strains used in this study.

| **Strain** | **Relevant characteristics** | **Source / Reference** |
| --- | --- | --- |
| PP2M350 | Environmental isolate M2 carrying pRGPdCas9bad; Kan^R^ | This study |
| PP2M353 | Environmental isolate M2 carrying pRGPdCas9bad-*argH*; Kan^R^ | This study |
| PP2M354 | Environmental isolate M2 carrying pRGPdCas9bad-*eda*; Kan^R^ | This study |
| PP2M355 | Environmental isolate M2 carrying pRGPdCas9bad-*edd*; Kan^R^ | This study |
| PP2M357 | Environmental isolate M2 carrying pRGPdCas9bad-*pheA*; Kan^R^ | This study |
| PP2M358 | Environmental isolate M2 carrying pRGPdCas9bad-*pyrF*; Kan^R^ | This study |
| PP2M402 | Environmental isolate M2 carrying pRGPspdCas9bad; Kan^R^ | This study |
| PP2M404 | Environmental isolate M2 carrying pRGPspdCas9bad-*argH*; Kan^R^ | This study |
| PP2M405 | Environmental isolate M2 carrying pRGPspdCas9bad-*eda*; Kan^R^ | This study |
| PP2M406 | Environmental isolate M2 carrying pRGPspdCas9bad-*edd*; Kan^R^ | This study |
| PP2M408 | Environmental isolate M2 carrying pRGPspdCas9bad-*pheA*; Kan^R^ | This study |
| PP2M409 | Environmental isolate M2 carrying pRGPspdCas9bad-*pyrF*; Kan^R^ | This study |
| PP2M411 | Environmental isolate M2 carrying pRGPspdCas9bad-*gcd*; Kan^R^ | This study |
| PP2M413 | Environmental isolate M2 carrying pRGPspdCas9bad-*xylB*; Kan^R^ | This study |
| PP2M493 | Environmental isolate M2 carrying pRGPspdCas9bad-*ara dehydrogenase*; Kan^R^ | This study |
| PP2M494 | Environmental isolate M2 carrying pRGPspdCas9bad-*ara MFS permease*; Kan^R^ | This study |
| PP2M496 | Environmental isolate M2 carrying pRGPspdCas9bad-*ara permease*; Kan^R^ | This study |
| PP2M498 | Environmental isolate M2 carrying pRGPspdCas9bad-*ara regulator*; Kan^R^ | This study |
| PP2M500 | Environmental isolate M2 carrying pRGPspdCas9bad-*araA*; Kan^R^ | This study |
| PP2M502 | Environmental isolate M2 carrying pRGPspdCas9bad-*araD*; Kan^R^ | This study |
| PP2M503 | Environmental isolate M2 carrying pRGPspdCas9bad-*araX*; Kan^R^ | This study |
| PP2M504 | Environmental isolate M2 carrying pRGPspdCas9bad-*xyl acetyltransferase*; Kan^R^ | This study |
| PP2M506 | Environmental isolate M2 carrying pRGPspdCas9bad-*xyl dehydrogenase*; Kan^R^ | This study |
| PP2M508 | Environmental isolate M2 carrying pRGPspdCas9bad-*xyl permease*; Kan^R^ | This study |
| PP2M510 | Environmental isolate M2 carrying pRGPspdCas9bad-*xyl regulator*; Kan^R^ | This study |
| PP2M512 | Environmental isolate M2 carrying pRGPspdCas9bad-*xylonate transporter*; Kan^R^ | This study |
| PP2M514 | Environmental isolate M2 carrying pRGPspdCas9bad-*xylA*; Kan^R^ | This study |
| PP2M516 | Environmental isolate M2 carrying pRGPspdCas9bad-*xylD*; Kan^R^ | This study |
| PP2M518 | Environmental isolate M2 carrying pRGPspdCas9bad-*xylX*; Kan^R^ | This study |

**Table S7.** CRISPRi plasmids used in this study.

| **Plasmid(*)** | **Relevant characteristics** | **Source / Reference** |
| --- | --- | --- |
| pRGPdCas9bad | dual-inducible CRISPRi system carrying two repressor genes (*lacI, araC*) and a gene encoding the dCas9 protein from *S. pasteurianus* under the control of IPTG-inducible P*_lac_* promoter and a cloning site for sgRNA cloning under the control of arabinose-inducible P*_aracbad_* promoter; Kan^R^ | Gauttam et al., 2021 |
| pRGPspdCas9bad | dual-inducible CRISPRi system carrying two repressor genes (*lacI, araC*) and a gene encoding the dCas9 protein from *Streptococcus pyogenes* under the control of IPTG-inducible P*_tac_* promoter and a cloning site for sgRNA cloning under the control of arabinose-inducible P*_araCbad_* promoter; Kan^R^ | Gauttam et al., 2021 |
| pRGPdCas9bad-*argH* M2 | pRGPdCas9bad carrying the sgRNA coding sequence targeting *argH*; Kan^R^ | This study |
| pRGPdCas9bad-*eda* M2 | pRGPdCas9bad carrying the sgRNA coding sequence targeting *eda*; Kan^R^ | This study |
| pRGPdCas9bad-*edd* M2 | pRGPdCas9bad carrying the sgRNA coding sequence targeting *edd*; Kan^R^ | This study |
| pRGPdCas9bad-*pheA* M2 | pRGPdCas9bad carrying the sgRNA coding sequence targeting *pheA*; Kan^R^ | This study |
| pRGPdCas9bad-*pyrF* M2 | pRGPdCas9bad carrying the sgRNA coding sequence targeting *pyrF*; Kan^R^ | This study |
| pRGPspdCas9bad-*argH* M2 | pRGPspdCas9bad carrying the sgRNA coding sequence targeting *argH*; Kan^R^ | This study |
| pRGPspdCas9bad-*eda* M2 | pRGPspdCas9bad carrying the sgRNA coding sequence targeting *eda*; Kan^R^ | This study |
| pRGPspdCas9bad-*edd* M2 | pRGPspdCas9bad carrying the sgRNA coding sequence targeting *edd*; Kan^R^ | This study |
| pRGPspdCas9bad-*pheA* M2 | pRGPspdCas9bad carrying the sgRNA coding sequence targeting *pheA*; Kan^R^ | This study |
| pRGPspdCas9bad-*pyrF* M2 | pRGPspdCas9bad carrying the sgRNA coding sequence targeting *pyrF*; Kan^R^ | This study |
| pRGPspdCas9bad-*gcdF* M2 | pRGPspdCas9bad carrying the sgRNA coding sequence targeting *gcd*; Kan^R^ | This study |
| pRGPspdCas9bad-*xylB* M2 | pRGPspdCas9bad carrying the sgRNA coding sequence targeting *xylB*; Kan^R^ | This study |
| pRGPspdCas9bad-*ara dehydrogenase* M2 | pRGPspdCas9bad carrying the sgRNA coding sequence targeting *ara dehydrogenase*; Kan^R^ | This study |
| pRGPspdCas9bad-*ara MFS permease* M2 | pRGPspdCas9bad carrying the sgRNA coding sequence targeting *ara MFS permease*; Kan^R^ | This study |
| pRGPspdCas9bad-*ara permease* M2 | pRGPspdCas9bad carrying the sgRNA coding sequence targeting *ara permease*; Kan^R^ | This study |
| pRGPspdCas9bad-*ara regulator* M2 | pRGPspdCas9bad carrying the sgRNA coding sequence targeting *ara regulator*; Kan^R^ | This study |
| pRGPspdCas9bad-*araA* M2 | pRGPspdCas9bad carrying the sgRNA coding sequence targeting *araA*; Kan^R^ | This study |
| pRGPspdCas9bad-*araD* M2 | pRGPspdCas9bad carrying the sgRNA coding sequence targeting *araD*; Kan^R^ | This study |
| pRGPspdCas9bad-*araX* M2 | pRGPspdCas9bad carrying the sgRNA coding sequence targeting *araX*; Kan^R^ | This study |
| pRGPspdCas9bad-*xyl acetyltransferase* M2 | pRGPspdCas9bad carrying the sgRNA coding sequence targeting *xyl acetyltransferase*; Kan^R^ | This study |
| pRGPspdCas9bad-*xyl dehydrogenase* M2 | pRGPspdCas9bad carrying the sgRNA coding sequence targeting *xyl dehydrogenase*; Kan^R^ | This study |
| pRGPspdCas9bad-*xyl permease* M2 | pRGPspdCas9bad carrying the sgRNA coding sequence targeting *xyl permease*; Kan^R^ | This study |
| pRGPspdCas9bad-*xyl regulator* M2 | pRGPspdCas9bad carrying the sgRNA coding sequence targeting *xyl regulator*; Kan^R^ | This study |
| pRGPspdCas9bad-*xylonate transporter* M2 | pRGPspdCas9bad carrying the sgRNA coding sequence targeting *xylonate transporter*; Kan^R^ | This study |
| pRGPspdCas9bad-*xylA* M2 | pRGPspdCas9bad carrying the sgRNA coding sequence targeting *xylA*; Kan^R^ | This study |
| pRGPspdCas9bad-*xylD* M2 | pRGPspdCas9bad carrying the sgRNA coding sequence targeting *xylD*; Kan^R^ | This study |
| pRGPspdCas9bad-*xylX* M2 | pRGPspdCas9bad carrying the sgRNA coding sequence targeting *xylX*; Kan^R^ | This study |

**Table S8.** Oligonucleotides used in this study for creating CRISPRi plasmids.

| **Oligonucleotides** | **Sequence (5’ → 3’)** | **Plasmid construction/ purpose** |
| --- | --- | --- |
| **Oligonucleotides used for sgRNA cloning into *S. pasteurianus* dCas9 based CRISPRi vector (pRGPdCas9bad)** | | |
| pdCas9 sgRNA-rev (M2) | tatgcataataatgcattaaggacactgtatctgc | Standard reverse primer for sgRNA cloning in p  RGPdCas9bad |
| argH M2 sgRNA-fwd | CATGATCATGAATCGACCGAGGCGGTGAAGCgtttttgtactcgaaagagcctacaaaga | pRGPdCas9bad-*argH* M2 |
| eda M2 sgRNA-fwd | CATGATCATGAGATATCTTCCTCACGGGCGAgtttttgtactcgaaagagcctacaaaga | pRGPdCas9bad-*eda* M2 |
| edd M2 sgRNA-fwd | CATGATCATGATGGGTCCGTCGCTGGCCGCGgtttttgtactcgaaagagcctacaaaga | pRGPdCas9bad-*edd* M2 |
| pheA M2 sgRNA-fwd | CATGATCATGAACCGGCTGCTCGCCTTCGGCgtttttgtactcgaaagagcctacaaaga | pRGPdCas9bad-*pheA* M2 |
| pyrF M2 sgRNA-fwd | CATGATCATGAGTCACGGGAGGGGAAATCCAgtttttgtactcgaaagagcctacaaaga | pRGPdCas9bad-*pyrF* M2 |
| **Oligonucleotides used for sgRNA cloning into *S. pyogenes* dCas9 based CRISPRi vector (pRGPspdCas9bad)** | | |
| pydCas9 sgRNA-rev | agcagtaaaggtaccacgacagatagatctaaggaaaaagtttcgtgtgcttcg | Common reverse primer for sgRNA cloning in pRGPspdCas9bad |
| argH M2 sgRNA-fwd1 | cctaggatccCCAGGACTGATTGGTCTTGTgttttagagctagaaatagcaagttaaaat | pRGPspdCas9bad-*argH* M2 |
| eda M2 sgRNA-fwd1 | cctaggatccGCAGAGGCTGGGGGCGTTCAgttttagagctagaaatagcaagttaaaat | pRGPspdCas9bad-*eda* M2 |
| edd M2 sgRNA-fwd1 | cctaggatccGGGTGACCTCAAGGATGCGCgttttagagctagaaatagcaagttaaaat | pRGPspdCas9bad-*edd* M2 |
| pheA M2 sgRNA-fwd1 | cctaggatccCAGCGCCTTGAGCTCATGTTgttttagagctagaaatagcaagttaaaat | pRGPspdCas9bad-*pheA* M2 |
| pyrF M2 sgRNA-fwd1 | cctaggatccGATCAGGGGCGTCTGGCAGGgttttagagctagaaatagcaagttaaaat | pRGPspdCas9bad-*pyrF* M2 |
| gcd M2 sgRNA-fwd | cctaggatccTAGCCAGCGGCTTCCATTCTgttttagagctagaaatagcaagttaaaat | pRGPspdCas9bad-*gcdF* M2 |
| xylB M2 sgRNA-fwd | cctaggatccGGTAATGCGTATCGGTATGAgttttagagctagaaatagcaagttaaaat | pRGPspdCas9bad-*xylB* M2 |
| Ara dehydrogenase M2 sgRNA-fwd | cctaggatccTTGCATGCAACCTCCTCAGTgttttagagctagaaatagcaagttaaaat | pRGPspdCas9bad-*ara dehydrogenase* M2 |
| Ara MFS permease M2 sgRNA-fwd | cctaggatccAACCTCTTTTTGTTGTTTTAgttttagagctagaaatagcaagttaaaat | pRGPspdCas9bad-*ara MFS permease* M2 |
| Ara permease M2 sgRNA-fwd | cctaggatccGAGCCGCTGCTCCTGAATCAgttttagagctagaaatagcaagttaaaat | pRGPspdCas9bad-*ara permease* M2 |
| Ara regulator M2 sgRNA-fwd | cctaggatccCGGTAGTTCATGGGGGGCGCgttttagagctagaaatagcaagttaaaat | pRGPspdCas9bad-*ara regulator* M2 |
| araA M2 sgRNA-fwd | cctaggatccACTCCTGTAACAAAGAAAGAgttttagagctagaaatagcaagttaaaat | pRGPspdCas9bad-*araA* M2 |
| araD M2 sgRNA-fwd | cctaggatccGTTTGTTATCAGACATAAGCgttttagagctagaaatagcaagttaaaat | pRGPspdCas9bad-*araD* M2 |
| araX M2 sgRNA-fwd | cctaggatccGCATGCTTGTGCTCGCTTGTgttttagagctagaaatagcaagttaaaat | pRGPspdCas9bad-*araX* M2 |
| Xyl acetyltransferase M2 sgRNA-fwd | cctaggatccGGCTGCATCTCTGCTCGTGGgttttagagctagaaatagcaagttaaaat | pRGPspdCas9bad-*xyl acetyltransferase* M2 |
| Xyl dehydrogenase M2 sgRNA-fwd | cctaggatccCTCCAGCTTCAACTGCGTGCgttttagagctagaaatagcaagttaaaat | pRGPspdCas9bad-*xyl dehydrogenase* M2 |
| Xyl permease M2 sgRNA-fwd | cctaggatccATGGTCCATAACCAGCCTCTgttttagagctagaaatagcaagttaaaat | pRGPspdCas9bad-*xyl permease* M2 |
| Xyl regulator M2 sgRNA-fwd | cctaggatccGTCCGACATGAATCCACCTTgttttagagctagaaatagcaagttaaaat | pRGPspdCas9bad-*xyl regulator* M2 |
| Xylonate transporter M2 sgRNA-fwd | cctaggatccGGGCACTGTGTCCTTGGCTTgttttagagctagaaatagcaagttaaaat | pRGPspdCas9bad-*xylonate transporter* M2 |
| xylA M2 sgRNA-fwd | cctaggatccCCTGCAAATGAAGTCATGAAgttttagagctagaaatagcaagttaaaat | pRGPspdCas9bad-*xylA* M2 |
| xylD M2 sgRNA-fwd | cctaggatccGACTGCGCAGCGGGCGTTTCgttttagagctagaaatagcaagttaaaat | pRGPspdCas9bad-*xylD* M2 |
| xylX M2 sgRNA-fwd | cctaggatccGCGTTATCGGCATGGCAGCTgttttagagctagaaatagcaagttaaaat | pRGPspdCas9bad-*xylX* M2 |

**Table S9.** The details of the putative essential genes for *P. putida* M2 whose expression has been downregulated in this study along with the target sequence for creating corresponding sgRNA for representative genes in pentose sugar utilization.

| **Target gene** | **Strains** | **Target sequence** |
| --- | --- | --- |
| Argininosuccinate lyase (*argH*) | PP2M353 | ATCGACCGAGGCGGTGAAG |
|  | PP2M404 | CCAGGACTGATTGGTCTTGT |
| 2-keto-3-deoxy-6phosphogluconate aldolase (*eda*) | PP2M354 | GATATCTTCCTCACGGGCGA |
|  | PP2M405 | GCAGAGGCTGGGGGCGTTCA |
| Phosphogluconate dehydratase (*edd*) | PP2M355 | TGGGTCCGTCGCTGGCCGCG |
|  | PP2M406 | GGGTGACCTCAAGGATGCGC |
| Chorismite mutase (*pheA*) | PP2M357 | ACCGGCTGCTCGCCTTCGGC |
|  | PP2M408 | CAGCGCCTTGAGCTCATGTT |
| Orotidine-5’-phosphate decarboxylase (*pyrF*) | PP2M358 | GTCACGGGAGGGGAAATCCA |
|  | PP2M409 | GATCAGGGGCGTCTGGCAGG |
| *gcd* | PP2M411 | TAGCCAGCGGCTTCCATTCT |
| *xylB* | PP2M413 | GGTAATGCGTATCGGTATGA |
| *ara dehydrogenase* | PP2M493 | TTGCATGCAACCTCCTCAGT |
| *ara MFS permease* | PP2M494 | AACCTCTTTTTGTTGTTTTA |
| *ara permease* | PP2M496 | GAGCCGCTGCTCCTGAATCA |
| *ara regulator* | PP2M498 | CGGTAGTTCATGGGGGGCGC |
| *araA* | PP2M500 | ACTCCTGTAACAAAGAAAGA |
| *araD* | PP2M502 | GTTTGTTATCAGACATAAGC |
| *araX* | PP2M503 | GCATGCTTGTGCTCGCTTGT |
| *xyl acetyltransferase* | PP2M504 | GGCTGCATCTCTGCTCGTGG |
| *xyl dehydrogenase* | PP2M506 | CTCCAGCTTCAACTGCGTGC |
| *xyl permease* | PP2M508 | ATGGTCCATAACCAGCCTCT |
| *xyl regulator* | PP2M510 | GTCCGACATGAATCCACCTT |
| *xylonate transporter* | PP2M512 | GGGCACTGTGTCCTTGGCTT |
| *xylA* | PP2M514 | CCTGCAAATGAAGTCATGAA |
| *xylD* | PP2M516 | GACTGCGCAGCGGGCGTTTC |
| *xylX* | PP2M518 | GCGTTATCGGCATGGCAGCT |

**Table S10.** Strains used in this study.

| **Strain (*)** | **Relevant characteristics** | **Source / Reference** |
| --- | --- | --- |
| **Strains** | | |
| *E. coli* DH5α | F^-^ *φ*80*lacZ*∆M15 ∆(*lacZYA-argF*) U169 *endA1 recA1 hsdR17* (r_k_^-^, m_k_^+^) *supE44 thi^-1^ gyrA996 relA1 phoA* | Hanahan (1983) |
| *P. putida* KT2440 | Wild type | ATCC 12633 |
| *P. putida* M2 | Accession number JADOUD010000001 | This study |
| *P. putida* PP2M403 (JPUB_020087) | *P. putida* M2 carrying pRGPspdCas9bad-*argB* M2; Kan^R^ | This study |
| *P. putida* PP2M404 (JPUB_020109) | *P. putida* M2 carrying pRGPspdCas9bad-*argH* M2; Kan^R^ | This study |
| *P. putida* PP2M405 (JPUB_020111) | *P. putida* M2 carrying pRGPspdCas9bad-*eda* M2; Kan^R^ | This study |
| *P. putida* PP2M406 (JPUB_020113) | *P. putida* M2 carrying pRGPspdCas9bad-*edd* M2; Kan^R^ | This study |
| *P. putida* PP2M408 (JPUB_020117) | *P. putida* M2 carrying pRGPspdCas9bad-*pheA* M2; Kan^R^ | This study |
| *P. putida* PP2M409 (JPUB_020119) | *P. putida* M2 carrying pRGPspdCas9bad-*pyrF* M2; Kan^R^ | This study |
| *P. putida* PP2M491 | *P. putida* M2 carrying pRGPspdCas9bad; Kan^R^ | This study |
| *P. putida* PP2M512 (JPUB_020163) | *P. putida* M2 carrying pRGPspdCas9bad-*xyl transporter* M2; Kan^R^ | This study |
| *P. putida* PP2M516 (JPUB_020171) | *P. putida* M2 carrying pRGPspdCas9bad-*xylD* M2; Kan^R^ | This study |
